# Unsupervised decomposition of natural monkey behavior into a sequence of motion motifs

**DOI:** 10.1101/2023.03.04.531044

**Authors:** Koki Mimura, Jumpei Matsumoto, Daichi Mochihashi, Tomoaki Nakamura, Toshiyuki Hirabayashi, Makoto Higuchi, Takafumi Minamimoto

## Abstract

Nonhuman primates (NHPs) exhibit complex and diverse behavior that typifies advanced cognitive function and social communication, but quantitative and systematical measure of this natural nonverbal processing has been a technical challenge. Specifically, a method is required to automatically segment time series of behavior into elemental motion motifs, much like finding meaningful words in character strings. Here, we propose a solution called SyntacticMotionParser (SMP), a general-purpose unsupervised behavior parsing algorithm using a non-parametric Bayesian model. Using three-dimensional posture-tracking data from NHPs, SMP automatically outputs an optimized sequence of latent motion motifs classified into the most likely number of states. When applied to behavioral datasets from common marmosets and rhesus monkeys, SMP outperformed conventional posture-clustering models and detected a set of behavioral ethograms from publicly available data. SMP also quantified and visualized the behavioral effects of chemogenetic neural manipulations. SMP thus has the potential to dramatically improve our understanding of natural NHP behavior in a variety of contexts.

## Introduction

In humans and other primates, complex and dynamical behavioral sequences consisting of gaze, facial expressions, postures, and body movements serve as expressions of internal states such as emotion and intention (i.e. nonverbal expression), which are fundamental for normal social life(1)(2). Nonhuman primates (NHPs), such as macaque and marmoset monkeys, have been shown to interact socially through such nonverbal expressions while they perform higher cognitive/motor functions, thus providing unique opportunities for modeling human brain function in healthy and diseased contexts(3)(4). A variety of NHP models of dysfunction in emotion and social communication have been proposed and studied using pharmacological interventions and genetic modifications at the adult or developmental stages(5)(6)(7)(8). In addition, recent advances in genetic manipulation techniques, such as chemogenetics, have allowed reversible manipulation of activity in specific brain circuits of freely moving monkeys, opening up valuable avenues for understanding the neural mechanisms that govern internal states(9)(10)(11). However, previous NHP studies have focused on measuring behavioral indicators based on the experimenters’ hypotheses, running the risk of overlooking changes in animal behavior that are beyond the scope of the prediction. Thus, advancing our understanding of the brain mechanisms that underlie internal states requires a quantitative, data-driven recapture of natural NHP behavior(12)(13); however, the lack of a method for doing this has created a bottleneck in this research field.

Recent developments in video-based motion-tracking systems have enabled the automated acquisition of large-scale behavioral data(9)(14)(15)(16). These data can be analyzed by machine learning to automatically segment and extract recurrent behaviors (ethograms) from the data, thus replacing human observation. Programmatically, this means automatically determining the starting and ending points of all data segments corresponding to a given ethogram, even for non-predetermined behavior. Several studies have proposed ethogram-detection algorithms, but each has certain limitations. Ballesta et al. (2014) proposed a method for automatically detecting ethograms(17); however, it requires manually set *ad hoc* extraction parameters for the predetermined target ethograms, making it unsuitable as an objective means of evaluating natural NHP behavior. Other studies have automatically classified NHP behavior by using a simple clustering method that has been successfully applied to rodents and insects to look at NHP body postures that correspond to different ethograms(14)(15). This method may be useful for studying NHPs in a confined range of research interests, such as when all postures in the data can be completely mapped to a set of ethograms of interest. However, comprehensive NHP behavioral analysis must target a broad range of ethograms that are beyond the scope of these posture-based classification methods—including temporally dynamic changes in multiple postures (e.g., jumps, turns, and catching prey), as well as complex and abstract changes, such as those that occur during social play(18)(19)(20).

Another framework for unsupervised behavioral data segmentation is one that fits a generative model to time series of data. In this scenario, observed behavioral parameters are modeled as probabilistic implements of a sequence of categorical and reproducible latent states (motion motifs)(21)(22). This framework has shown promise in analyzing behavioral data from rodents, fish, and insects for identifying novel ways of capturing prey(23), sorting neuropharmacological effects(24), and mapping animal internal states by simultaneously recording neural activity(25)(26)(27). This technique is also gaining prominence in the fields of computer vision and human robotics as a way to computationally estimate the meaning of behavior(28)(29)(30). However, to the best of our knowledge, such an algorithm has not been verified as available for NHPs, likely because the current algorithms were not developed or optimized to target the complexity and diversity of NHP behavioral ethograms. In addition, even if the current algorithms were available for NHPs, they remain somewhat subjective; for unsupervised behavior segmentation, the determination of the number of latent classes (class size) has a significant impact on the results, but has usually been left to the subjective judgment of the researchers(14)(31). Ideally, the optimal class size should also be determined in an unsupervised manner(32)(33).

Here, we propose a new framework called SyntacticMotionParser (SMP), which allows for quantitative and automatic parsing of NHP behavior into a set of motion motifs. Using SMP, three-dimensional (3D) motion-tracking data can be described as a stochastic generative process of motion motifs, each of which is optimally segmented by unsupervised machine learning (Fig. 1a). Specifically, the internal variability of individual motifs is regressed by a Gaussian process (GP) and simultaneously clustered by its hyperparameters, while the number of motif classes is automatically optimized by a Bayesian nonparametric model with hierarchical Dirichlet process (HDP) (Fig. 1b). We demonstrate that SMP can characterize and describe different styles of common marmoset (*Callithrix jacchus*) feeding behavior. SMP was also able to extract several ethograms unique to rhesus monkeys (*Macaca mulatta*) from publicly available motion-tracking data and describe their patterns and temporal sequences(14). Critically, SMP was also able to automatically detect and describe the changes in marmoset behavior that resulted from chemo-genetic manipulation of specific neural circuits(9), without any prior information.

**Fig. 1.**
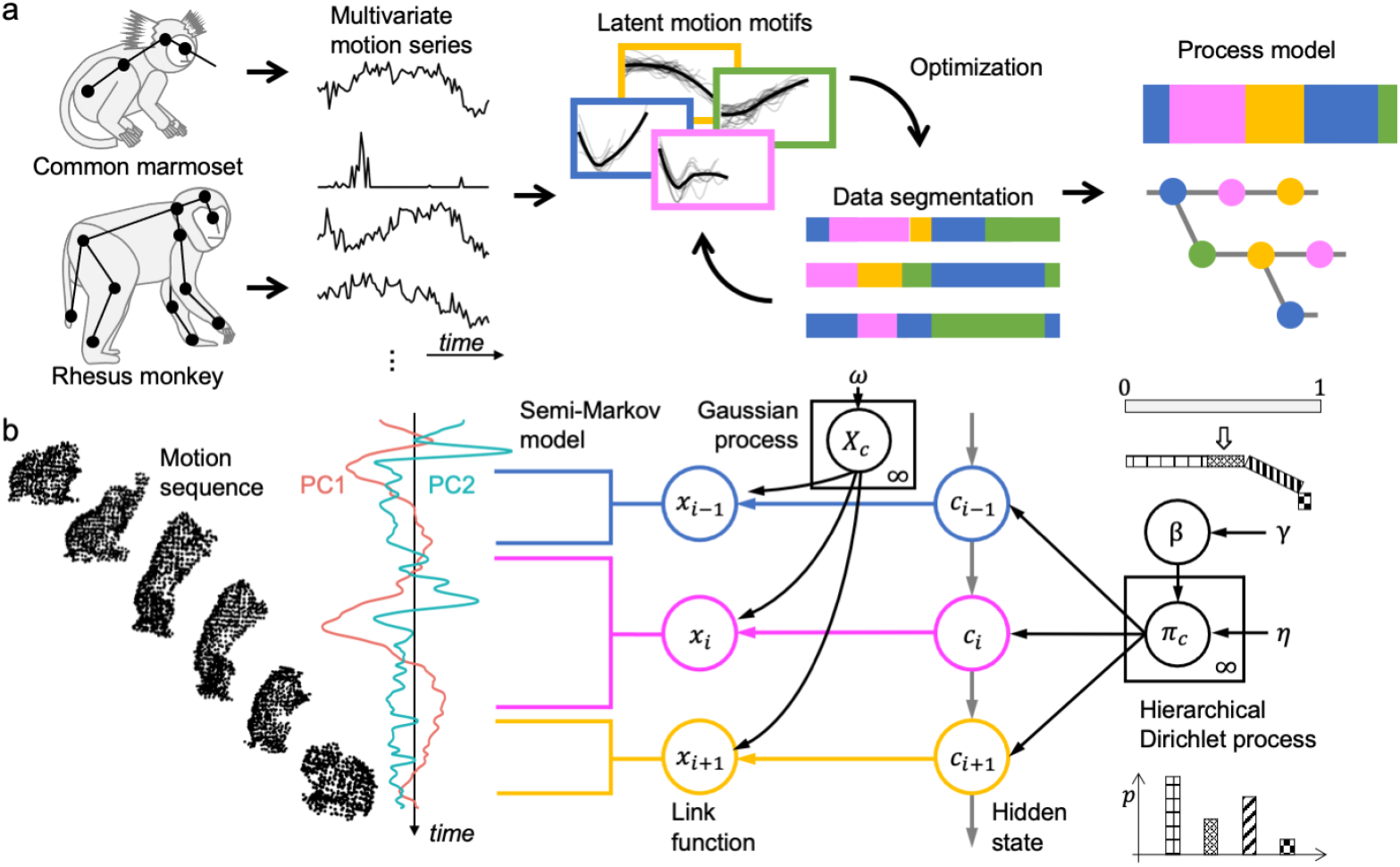
Dynamic temporal segmentation of NHP motion time series using a generative model. **(a)** Illustration of the generative description workflow of NHP behavior. The input behavior series is segmented into a set of latent motion motifs, which are partially reproducible trends of multivariate motion parameters. The properties of motion motifs and their probabilistic interconnectedness, i.e., “the grammar of behavior,” are optimized by data-driven machine learning and generatively describe behavior as a stochastic process model. **(b)** A graphical model of dynamical decomposition of monkey behavior using the nonparametric Bayesian model, SMP. The two major principal component scores (PC1 and PC2) extracted from the multivariate time series of monkey behavior were optimally segmented by this model into a time series of discretely labeled hidden state c, using GP as the nonlinear regression link function, a hidden semi-Markov model for estimating the temporal breakpoints, and HDP including the stick-breaking process for estimating state class size. NHP, nonhuman primate; SMP, SyntacticMotionParser; GP, Gaussian process; HDP, hierarchical Dirichlet process.

## Results

### Unsupervised detection of latent motion motifs from free-feeding marmoset behavior

We first a ssessed the ability of SMP’s computational segmentation to identify changes in internal states. Internal states were inferred from simple goal-directed behaviors, which included stereotyped and reproducible motion series such as searching, discovering, approaching, and eating. We recorded free-feeding behavior of four adult marmosets, during which they fed wherever, whenever, and however they wanted (Fig. 2a-b, Supplementary Fig. 1). Using an original marker-less motion-tracking system(9)(34)(35), the marmoset behavior was semi-automatically captured as 3D trajectories of the four body parts (*Face, Head, Trunk*, and *Hip*; Fig. 2c-d and Supplementary Movie 1). For the SMP analysis, we manually extracted trajectory data (approx. 20 s) centered on the timing of feeding based on video clips (N = 59; Supplementary Table1). We assumed that this 20-s feeding behavior would consist of a few motion units, each lasting several seconds. We applied SMP to the 1st and 2nd principal component (PC) scores of the free-feeding data (51 × 20 s) including 3D trajectories of body parts (13 parameters)(Fig. 2e). SMP successfully segmented the data into about 430 motion motifs (433.8 ± 5.2, mean ± sd) across simulations with various initial class sizes (random seed, n = 23). In those simulations, the number of motif classes converged to a unimodal posterior distribution with a median value of 18, which indicates the optimal class size (Fig. 2f-g, red). Analysis of the seven simulations that yielded 18 motif classes revealed that the frequency distribution of the motifs was similar regardless of the initial class size (X-sq = 30.13, df = 102, p-value = 1.00, Pearson’s Chi-squared test, Supplementary Fig. 2). Figure 2h shows 18 motion motifs characterized by a distinct set of PC-score dynamics segmented by SMP with an initial class size of 22 (which was used in the subsequent analyses). These motion motifs were commonly observed in the four marmosets without significant individual unique distribution bias (X-sq = 62.99, df = 51, p-value = 0.121, Pearson’s Chi-squared test, Supplementary Table 2). Successful convergence of dynamic motion segmentation appeared to result from the combination of GP and HDP inherent in SMP; the GP works as a link function that represents dynamic postural changes, while the HDP optimizes the class size(36)(37)(38). To benchmark SMP performance, we compared it with the performance of other statistical models. When using GP as a link function, but not using HDP, the class size depended on the initial value and the optimal class size was not determined (Fig. 2f, Model1). Similarly, when using HDP with a different link function (autocorrelation and static; Fig. 2f, Models 2 and 3, respectively), the class size again depended on the initial value and no information about optimal class size was provided. Thus, SMP that incorporates both HDP and GP demonstrated a significant advantage in simultaneously providing flexible regression to explain free-feeding behavior and returning an optimal class size of motion motifs inherent in the data.

**Fig. 2.**
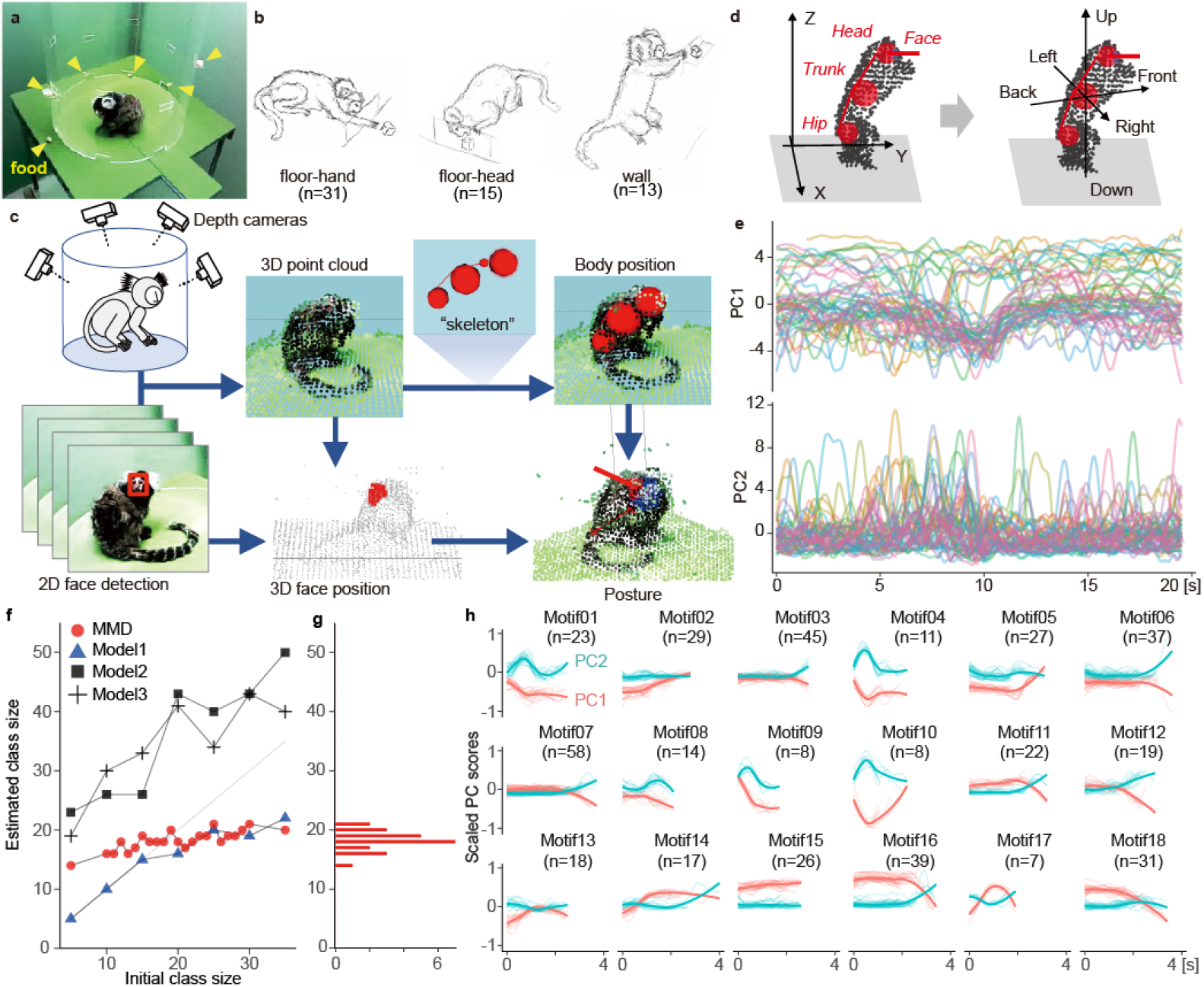
Motion tracking and computational segmentation in freely moving common marmosets. **(a)** Experimental setup for marmoset free-feeding behavior. Yellow arrowheads indicate the location of the food reward. **(b)** Illustrations showing the subtypes of feeding behavior: using hands to take food from the floor (*floor-hand*, left), using head and mouth directly (*floor-head*, middle), and taking food from the wall (*wall*, right). **(c)** Data-flow diagram of the marker-less motion-tracking system. The positions of marmoset body parts, *Head, Neck, Trunk*, and *Hip* were estimated by a skeleton model fitting. The *Face* position was also estimated by the projection of the face rectangle on RGB images to the point cloud. See also Supplementary Movie 1. **(d)** The posture parameters were transformed from distance-from-the center coordinates (left) to *Trunk*-centered coordinates (right). **(e)** Time series of the first and second principal component scores (PC1 and PC2) with 10 Hz resolution for a total of 1,020 s of data comprising 51 sets of 20-s data. **(f)** The class size of the elemental motion units during the free-feeding behavior was estimated by four statistical models: the proposed SMP method (HDP + GP, red), Model1 (GP, blue), Model2 (HDP + linear regression, square), and Model3 (HDP, cross). **(g)** Using MDD, the number of fragment classes converged to a unimodal posterior distribution with a median of 18. **(h)** Examples of a set of 18 motion units estimated by SMP simulation with 22 initial classes. SMP, SyntacticMotionParser; HDP, hierarchical Dirichlet process; GP, Gaussian process.

### SMP can describe and reproduce sequences of motion motifs from goal-directed marmoset behavior

Having shown that SMP performed consistently in unsupervised segmentation of marmoset free-feeding behavior, we next investigated how feeding behavior can be described with a sequence of motion motifs detected by SMP. Figure 3a presents a 20-s data sample that included two feedings. SMP decomposed these data into a sequence of eight motion motifs from four classes. Because each motion motif represents typical body motion trajectories that are transformed into temporal dynamics of two PCs (Fig. 2h), corresponding body motion parameters can be retrieved by inverse calculation from the mean value of the PC scores. For example, when a series of movements represented by motifs 2, 3, 7, and 8 were reconstructed in the order of observation, marmoset behavior was able to be retrieved as the dynamic relative positions of the *Face, Head, Trunk*, and *Hip* (Fig. 3d, Supplementary Movie 2), similar to those from the original data (Fig. 3b-c). When we attempted a similar reproduction of movements using a standard posture model (the 2D UMAP; uniform manifold approximation and projection)(18)(19) with the k-means clustering method, the motions were unnatural and resembled a stop-motion animation of static postures stitched together (Fig. 3e). With the same initial class size as that of the SMP (k = 18), the posture model decomposed the data into significantly shorter sub-second fragments (UMAP: 0.5 ± 1.16 s, median ± sd; SMP: 2.2 ± 0.59 s; BM statistic = -443.83, df = 1253.4, p-value < 2.2e-16) (Fig. 3f). These results demonstrate that SMP is unique in that it allows for a simple description of the multi-second dynamical structure observed during feeding behavior.

**Fig. 3.**
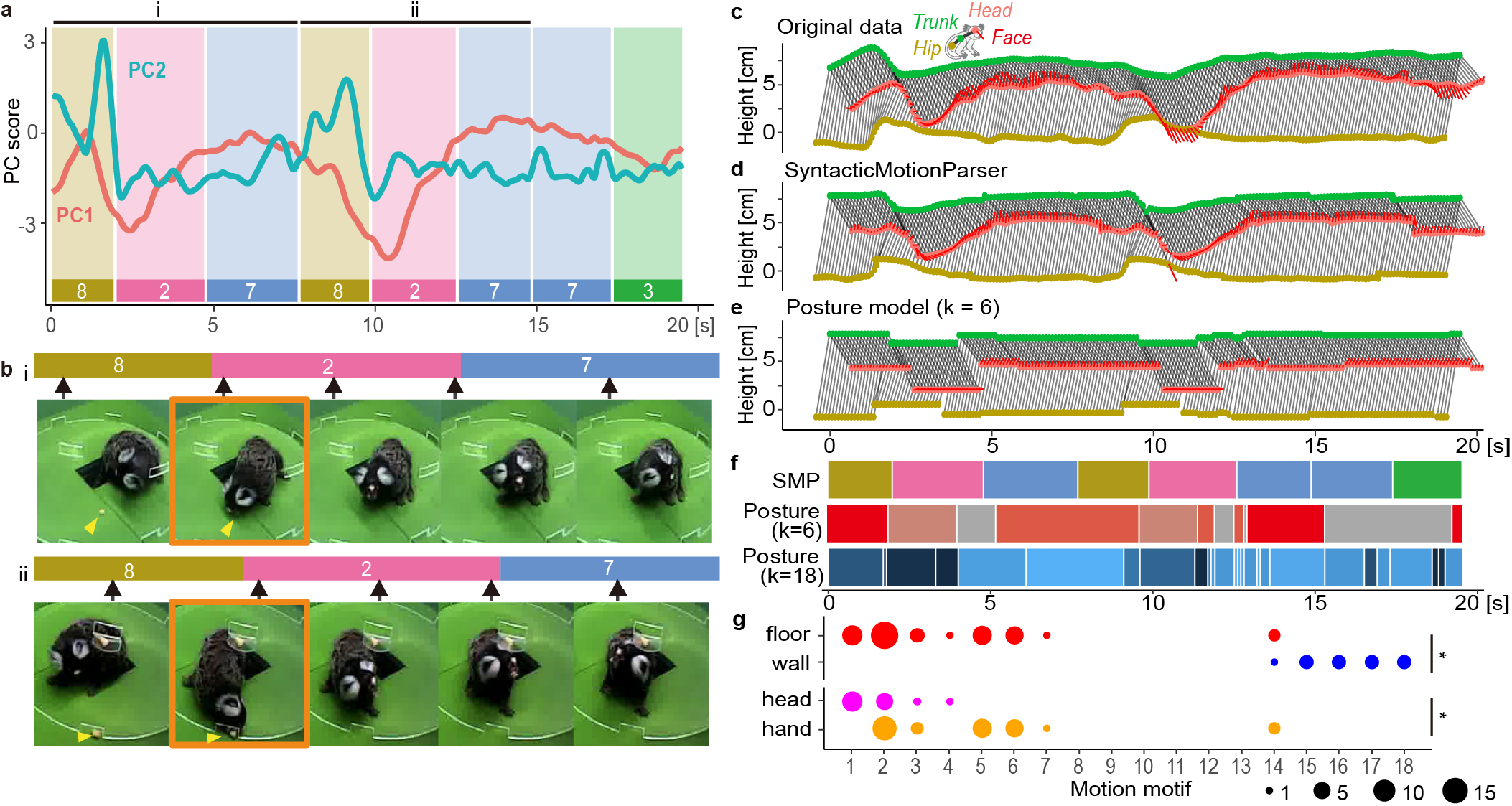
Example of marmoset behavior smoothly described by SMP-derived motion units. **(a)** Example of behavior segmentation. The curve trends represent the 1st and 2nd principal component scores (PC1 and PC2) of the motion parameters. The bottom lines and background rectangles represent the resulting eight segmented motion units with four types (8, 2, 7, and 3). **(b)** Two sequential images of marmoset free-feeding behaviors with three-segment arrays “8-2-7”, where the labels (i) and (ii) correspond to those in **a**. Black arrows indicate the position of the video frame in the segment represented by the upper bar. Yellow arrowheads indicate the location of the food reward. Orange rectangles indicate the images in which feeding occurred. **(c)** Time series of the original body movement trajectory during the data from **a**, with the X-axis representing time and the front-back direction, and the Y-axis representing the up-down direction. The *Face, Head, Trunk*, and *Hip* are color-coded in red, orange, green, and dark yellow, respectively. **(d)** The time series of the ideal body motion trajectory, which was pieced together from the eight motion units provided by SMP. **(e)** Time series connecting the six representative postures provided by the posture model, calculated by clustering the two-dimensional UMAP scores of all data using the k-means hierarchical clustering method (k = 6). See also Supplementary Movie 2. **(f)** Segmentation results of the motion sequence **c** using the proposed SMP method and the posture models (k = 6 and 18). **(g)** Dot plots showing the distribution of motion motifs in response to food placed on the wall and floor and in feeding approaches to food placed on the floor (*floor-hand* and *floor-head*). Size indicates the number of observations. *, p-value < 0.05 by Pearson’s Chi-squared test.

### SMP motion motifs quantitatively characterized classification by observation

As shown in Figure 3a, SMP described feeding behavior as a specific sequence motion motifs (e.g., 8–2–7). Because feeding behaviors can be classified manually into three subtypes according to food position and approach strategy (*floor-hand, floor-head*, and *wall*; Fig. 2b; Supplementary Table1), we next asked whether SMP-derived motion motifs would correspond to different feeding subtypes. Analysis revealed differences in motion motifs according to where the food was located (*wall* vs. *floor*), indicating that significantly different motifs were employed in the feeding segment (X-sq = 55.1, df = 11, p-value = 7.37e-08, Fig. 3g, top). We also found significant differences according to approach (*floor-hand* vs. *floor-head*), indicating that different motifs were employed depending on the approach strategy (X-sq = 22.6, df = 7, p-value = 1.97e-3, Fig. 3g, bottom). For comparison, we assessed the distribution of posture clusters during feeding using a conventional posture model with k = 18. Although the distributions of posture clusters at feeding times differed significantly between *wall* and *floor* (X-sq = 54.343, df = 10, p = 4.18e-08, Pearson’s Chi-squared test), they did not significantly differ between *floor-head* and *floor-hand* (X-sq = 4.7454, df = 8, p = 0.784, Pearson’s Chi-squared test; Supplementary Table 3). Thus, SMP could detect smaller, fine-grain differences in behavioral features than the conventional posture model could, meaning it has a higher behavioral resolution. Compared with existing posture-based methods, when given data that was manually and roughly clipped to the timing of specific events, SMP provided a description that better reflected the dynamics of the actual behavior.

### Using SMP to identify ethograms in freely moving macaques

Having demonstrated the effectiveness of SMP analysis of marmoset behavior during a goal-directed paradigm, we next applied SMP to data from freely moving NHPs to test whether it could extract what animals are doing (i.e., ethograms), in a data-driven manner. For this purpose, we used OpenMonkeyStudio, an open data source of macaque monkey behaviors(14), which consists of 3D trajectories of 13 key landmarks on a macaque body captured by multiple synchronized cameras (Fig. 4a, top). Similar to the data preprocessing in the marmoset analysis, the Front-Up coordinates were aligned to the *Head*-*Hip* axis (*Hip*-centered) to determine what each animal was doing rather than where it was (Fig. 4a, bottom). We assumed that the macaque behavior would comprise several kinds of motion motifs, each lasting around 15 s. The SMP segmented the data into about 230 motion motifs (234.2 ± 6.5, mean ± sd) in 16 random-seed simulations with various initial class sizes. The motifs comprised 10 different classes whose sizes did not vary depending on the initial values of the simulation (Fig. 4b). Figure 4c shows the PC waveforms of motion motifs that resulted from SMP extraction with an initial class size of 18. By inverse-calculating the postural parameters from the average trend of the PC waveforms, SMP resynthesized the ideal motion represented by each motion motif, as exemplified in Fig. 4d: walking (motif 1), climbing up and holding on upside down (motif 2), stepping down (motif 3), and climbing up and staying (motif 4), which are common motion ethograms of macaques. Furthermore, less common ethograms were also captured as independent classes, such as long-distance jumps along walls and to the floor (motif 8 and motif 9; Fig 4c, Supplementary Movie 3). These results demonstrate the versatility of SMP for detecting ethograms in freely moving NHPs from any dataset, with only a minimum assumption of their duration.

**Fig. 4.**
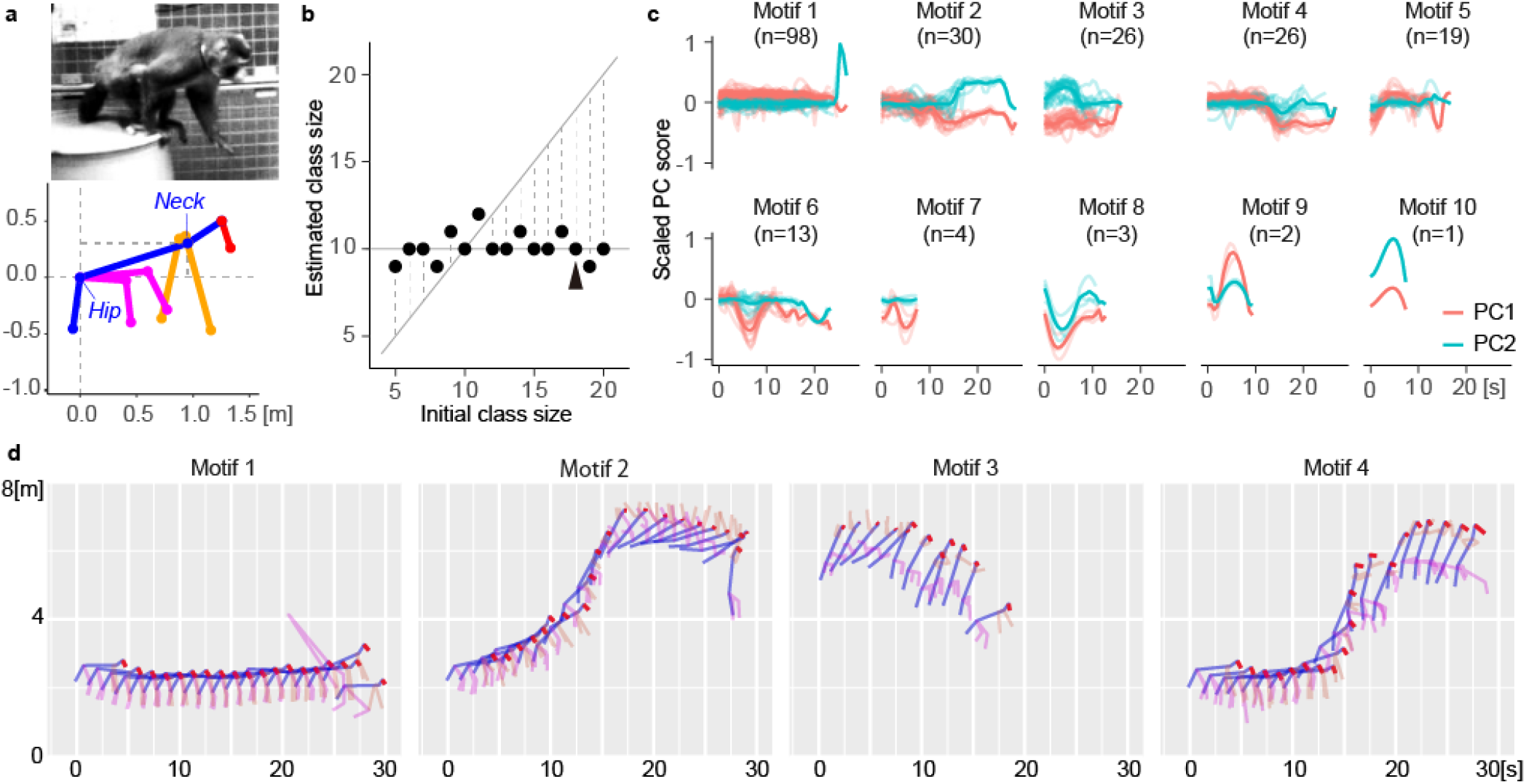
Independent macaque behavior was segmented into ethograms by SMP. **(a)** Published macaque image (top; adapted from OpenMonkeyStudio(14)) and corresponding posture keypoint coordinates, which have been rescaled as *Hip*-centered with the x-y plane including the *Hip*-*Neck* vector (bottom). **(b)** The median posterior distribution of class sizes was 10. **(c)** Example of the trends for the 1st and 2nd principal components (PC1 and PC2) of each motion unit with initial class size = 18 (arrowhead in **b**). **(d)** The synthesized ideal body trajectories in major motions are calculated by the inverse operation of PC analysis. Motion motifs 1–4 represent walking, climbing up and holding upside down, stepping down, and climbing up and staying, respectively (see also Supplementary Movie 3).

### Using SMP to characterize behavioral change induced by circuit manipulation

To identify how activity within a specific brain circuit can cause specific natural behaviors, we must be able to establish a method for unsupervised detection of behavioral changes. We therefore tested whether SMP could detect and characterize behavioral changes in free-moving marmosets that were induced by chemogenetic neural manipulation in our previous study(9). Marmosets that expressed the excitatory chemogenetic receptor hM3Dq in the unilateral substantia nigra (SN) spun themselves around in the direction contralateral to the activated SN, and this behavior began about 45 min after eating the chemogenetic activator (deschloroclozapine, DCZ) (Fig. 5a-b). Although this behavioral change can be quantified by counting the number of rotations, its characterization is not simple, as it is difficult to describe rotation behavior using conventional static postures in a series of snapshot images. We asked whether SMP could effectively capture this phenotype without any assumptions. We analyzed two sessions of 5-min motion-tracking data at four certain time windows (10-15, 35-40, 60-65, and 85-90 min) after DCZ or vehicle consumption. We used 3D trajectories of the *Head, Trunk*, and *Hip*, and their velocities in the postural coordinates where the *Head* was frontal and the *Trunk* was centered (Fig. 5c-d), and applied PC analysis to reduce their dimensions to two. With multiple initial class sizes, SMP consistently extracted seven classes of motion motifs (approx. 30-s) from the data (Fig. 5e). Temporal distribution of these motifs indicated that more than 90% of motifs 1 through 4 appeared later than 60 min after DCZ administration (Fig. 5f, red). Visualization of the reconstructed body movements through inverse PC analysis revealed that motion motifs 1-4 characterized the nature of the contralateral rotation with dynamical postural changes, which were clearly dissociable from normal directional changes (motifs 5 and 6 ipsilateral and contralateral, respectively) or exploratory behavior (motif 7) (Fig. 5g, Supplementary Movie 4). These results demonstrate that without any prior information, SMP could successfully capture and describe changes in unconstrained behavior that were induced by circuit manipulation. Importantly, it also indicated when and how the behavioral effects appeared, highlighting another advantage of this method in studying the causal relationship between natural behavior and brain function in NHPs.

**Fig. 5.**
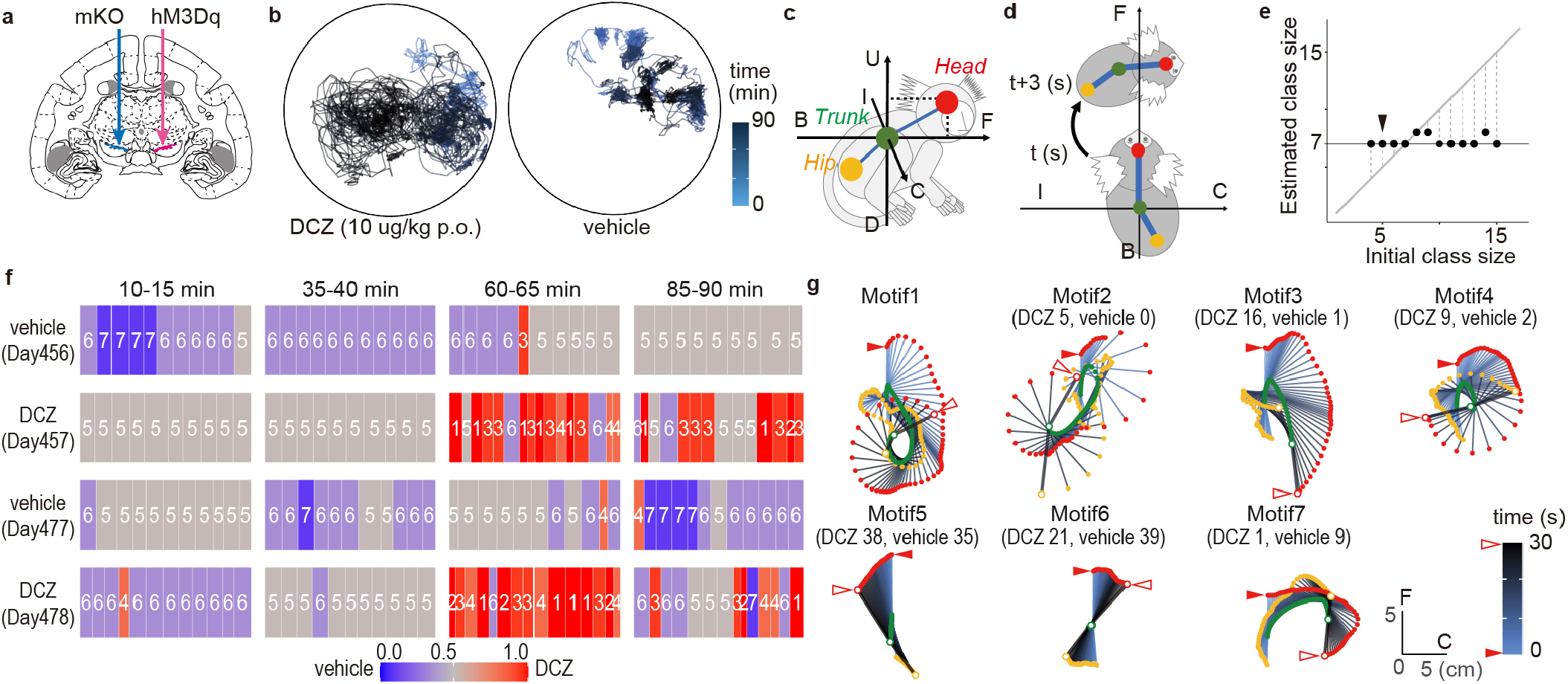
SMP characterized contralateral rotation behavior induced by chemogenetic neuro-manipulation. **(a)** Illustration of viral vector injection locations adapted from Mimura et al., 2021(9). AAV vectors expressing excitatory DREADD hM3Dq were injected into the unilateral substantia nigra (SN, red arrow). As a control, an AAV vector expressing a fluorescent marker (mKO) was injected into the contralateral SN (blue arrow). **(b)** Example of the top view of the *Head* trajectory of the marmoset after deschloroclozapine (DCZ; 10 μg/kg, per os (p.o.)) and vehicle administration. **(c)** Illustration of the posture coordinate. The horizontal mapping of the *Head* through the center of the *Trunk* was defined as the front-back (F-B) axis, and the axes orthogonal to the F-B axis horizontally and vertically were defined as the ipsilateral-contralateral (I-C) and vertical up-down (U-D) axes, respectively. **(d)** Body coordinates were used to determine relative positions every 3 s. **(e)** Estimation result of motion-motif class size. SMP was applied to four 5-min time windows each from two 90-min sessions after DCZ and vehicle p.o. The result was 7 kinds of motion motifs. **(f)** Example of a motion-motif sequence detected by SMP with initial class size = 5 (arrowhead in **e**). The time at which the motif was observed is indicated by tiles and color-coded by the percentage of occurrences in the DCZ data. The days since vector injection are shown below the dosing conditions. **(g)** The top view illustration of the body trajectories is represented by the motion motifs in **f**. Observed numbers in the DCZ and vehicle groups are under the motif numbers. The red and white arrowheads indicate the starting and ending points of the *Head* track, respectively. The *Head, Trunk*, and *Hip* positions are color-corded as **c**, with white at the ending points. The body positions are drawn at 0.5-s intervals with a gradation from blue to black from start to end (see also Supplementary Movie 4).

## Discussion

Here, we describe a novel computational framework, SMP, that quantitatively and automatically parses natural NHP behaviors into motion motifs, which can be used as meaningful metrics, like ethograms. As illustrated by its application to three NHP datasets in two monkey species, SMP is versatile and effective for quantifying and characterizing behaviors by simply adjusting the temporal window for motifs of interest, and not changing the model or process. SMP can also automatically detect and describe behavioral changes induced by neuronal manipulation. Thus, SMP can be widely applied for motion description and quantification of NHPs, offering the potential to dramatically improve our understanding of natural NHP behavior and the underlying brain functions in both normal and disease states.

Recent advances in machine learning allow for the collection of large-scale posture data, measured and estimated automatically from videos of freely moving NHPs(9)(14)(15)(16). At the same time, frame-by-frame classification of posture information can be used to detect ethograms from streaming behavior data(14). Although this type of classification has been shown to be effective in rodents(39), it may not be sufficient in NHPs because their natural ethograms are typically longer and more complex, which makes it more likely that the same postures occur in different contexts, as happens when climbing up and down. Indeed, in our demonstration, the conventional posture-classification method could not capture and reproduce NHP motion motifs because these methods transform inherently smooth and dynamic postural changes in a given ethogram into a constrained combination of static states, i.e., postures (Fig. 3e-f). In the case of rodents and insects, attempts have been made to address this issue by refining the conventional posture model to account for the semantic context(33)(40). The results are like stop-motion animations, forcing complex and expensive modifications to reproduce natural actions. Unlike these static posture models, our proposed SMP can provide dynamic and reproducible motion motifs—which are inherent in the behavior—as basic units. Using a probabilistic estimation of the type and occurrence of motifs through data-driven learning, SMP automatically parses streaming behavioral data into a series of motion motifs (or ethograms). This is similar to the way in which sentence structure analysis in natural language processing directly searches for meaningful morphemes like words, omitting detailed consideration of the smallest static unit, the character. SMP clearly distinguished climbing up from climbing down as different behavioral “words”, even though these two actions are indistinguishable in principle by posture alone (Fig. 4, Supplementary Movie 3). Thus, by estimating the grammar and syntax of NHP behavior, motion parsing might allow researchers to understand the internal state of animals or the functional significance of the behavior in terms of social communication and the underlying brain mechanisms, similar to what has already been shown to be effective for time-series correlation analysis with neural activity in rodents(24)(26).

Our results also demonstrate that SMP represents a major advance in how the behavioral effects of neural manipulation can be evaluated. While conventional methods have long relied on subjectively defined ethograms or task-controlled behavior, SMP employs automatic detection and description of changes in natural behavior, such as the frequency and type of motion motifs. Our demonstration showed that SMP can isolate and resynthesize abnormal contralateral turning that was triggered by chemogenetic neuronal manipulation, even under severe constraints such as limited tracking points that lacked facial coordinates (Fig. 5, Supplementary Movie 4). When chemogenetics is applied to NHP brain circuits that govern complex behaviors (e.g., the dorsolateral prefrontal cortex and caudate nucleus)(41), SMP may be able to detect resulting behavioral changes not only as abnormal motifs, but also as abnormalities in motion-motif sequences (e.g., “8-2-7” in feeding behavior in Fig. 3) or their transition structures. In addition, by building a large dataset of natural NHP behavior, we can address behavioral variations across individuals. Our results suggest that SMP holds enormous potential for quantifying and describing behavioral changes far beyond what can be measured by conventional observational or task-based methods.

SMP implements two extensions to its generative model, HPD and GP, which address the following two major issues related to statistical estimation of latent states. The first issue is determining how many latent states exist in the behavioral data, which is always a problem when labeling discrete data elements as “characters” or “words”. Because the class size of a latent state model is usually given as an *ad hoc* fixed parameter(42), researchers have struggled to determine which is the most likely result when running simulations under a variety of conditions, each returning a plausible result(14). The HDP is theoretically guaranteed to yield a single estimated class size for any initial value in an ideal situation with infinite iterations, which eliminates the need to select an optimal result(36)(37)(38). Our comparison of multiple initial values was not for a posteriori sorting as in previous studies(33), but rather it was to confirm that sufficient iterations were performed for convergence. In fact, all of our simulations returned a simple unimodal posterior distribution of the class sizes after 100 iterations (Fig. 2f-g, 4b, and 5e), from which the most likely class size was uniquely and automatically determined. The second issue is related to the link function of the latent state model. In a typical latent state model, each state is assumed to be static, as is the posture, which is why a link function is needed to regress and represent complex internal variability such as motion. We chose GP as the regression link function because it has been used in human behavior analyses(29)(30)(43), and because it allows for more flexible regression than conventional auto-regressive filtering or Bernoulli generalized linear models(25)(26).

By incorporating both GP and HDP, SMP is able to automatically estimate all parameters, except for the three hyperparameters and hidden state intervals (min, max, and mean). For the three hyperparameters, ω, γ, and η (parameters that control error coefficients, sparsity of the underlying state transition matrix, and expected number of different hidden states, respectively; Fig. 1b), we showed that the basic parameter settings are useful for NHP behavior quantification, and that the values were similar to those in our human motion analysis(29). The hyperparameters might need to be tuned for data with different variance assumptions. As for the state interval parameters, they are major adjustment factors that directly reflect the temporal extent of the motion motifs of interest. Although there are no theoretical limitations on the intervals, longer intervals increase computational cost and decrease the frequency of reproducible GP regression curves. Therefore, when trying different intervals, we recommend that the parameterized frame length first be fixed and the time resolution of the data be adjusted, as we demonstrated. While a similar latent model with HDP and GP was able to identify human motion as accurately as manual annotation(29), the current study is the first application of this type of model extension for exploratory behavior quantification in NHPs, for which the GP kernel function was optimized.

SMP will open the door to quantitative, rigorous, and comprehensive research of natural monkey behavior, which is needed in a wide range of scientific disciplines, including neuroscience, ethology, and developmental and evolutionary biology. Application of SMP also has a potential use in drug development and engineering brain-machine interfaces and other clinical devices. Neural recording and circuit manipulation of NHPs in free-moving conditions(9)(44) would bear real fruit when combined with computational behavioral description. SMP overcomes the motion annotation bottleneck of NHP experiments and automatically describes the diversity and complexity of the dynamical natural behavior and the behavioral changes associated with brain circuit manipulation, therefore increasing the value of NHPs in the fields of neuroscience and psychology by opening a new behavioral research field—natural nonverbal processing.

## Materials and methods

### Animals

Four laboratory-bread adult common marmosets were used (2 males, 2 females; 1.4–6.4 years old; 290–400 g, Supplementary Table1). Each cage was exposed to a 12/12-h light-dark cycle. Room temperature and humidity were maintained at 27–30°C and 40%–50%, respectively. All experimental procedures were performed in accordance with the Guide for the Care and Use of Nonhuman Primates in Neuroscience Research (Japan Neuroscience Society; https://www.jnss.org/en/animal_primates) and were approved by the Animal Ethics Committee of the National Institutes for Quantum Science and Technology (#11-1038).

### Behavior test

Behavior experiments were conducted in a sound-attenuated room (O’hara & Co., Ltd., Tokyo, Japan; 2.4 m (h) × 1.2 m (w) × 1.6 m (d)), which was apart from the colony room. Vocalizations from the colony room could not be detected in the experimental room. The temperature was maintained at 27–30°C and relative humidity was 30%–40%. The internal space of the sound-attenuated room was ventilated and illuminated with fluorescent lighting. The experiments were performed between 11:00 and 16:00.

Before the experimental sessions, each subject was transferred individually from the colony room to the experimental room in a small transport chamber (O’hara & Co., Ltd., Tokyo, Japan; 300 mm (h) × 100 mm (w) × 100 mm (d)). Once in the experimental room the transport chamber was placed under a table with a green top (O’hara & Co., Ltd., Tokyo, Japan; 0.5 m × 0.5 m × 0.5 m), upon which rested a cylindrical test chamber made from transparent acrylic (0.4 m (r) × 0.5 m (h)). The bottom of the test chamber had a door that when opened allowed the marmoset to enter from the transport chamber (Supplementary Fig. 1a). The test chamber had 16 feeding ports (30 mm × 15 mm) located at 45-degree intervals on the floor (8 ports) and on the wall at heights of 150 mm and 200 mm (4 each, alternately arranged). Platforms (30 mm × 15 mm) were set up on the outside of the wall ports where food rewards could be placed (Supplementary Fig. 1b). For 3D data acquisition, 4 depth cameras (RealSense Depth Camera R200, Intel, Santa Clara, USA) were placed around the chamber at 90-degree intervals (Supplementary Fig. 1b), and were connected in parallel to a PC (Windows 10, 64-bit) using USB-C cables (U3S1A01C12-050, Newnex Technology, Santa Clara, USA; the distance was 1-5 m). The subjects were allowed to adapt to the transport procedure and experimental environment for two consecutive days before behavioral testing.

Each behavioral test started when a marmoset entered the test chamber from the transport chamber through the floor entrance, and the recording of movements lasted until they finished eating all the food placed in the feeding ports, lasting up to 30 min. Subjects freely moved around the test chamber and, whenever they wanted, ate 7–9 pieces of sponge cake (approximately 2 g each); one cake was located on the near side of the entrance to entice them to enter the chamber, two on the wall ports, two on the near side of the floor ports, and the others 3 cm away from the floor ports (Fig. 2a, Supplementary Fig. 1 b). Observed feeding behaviors were manually classified into three subtypes: using hands to take food from the floor (*floor-hand*), using head and mouth directly (*floor-head*), and taking food from the wall (*wall*) (Fig. 2b, Supplementary Table1). Fifty-one sets of 20 s (10 s before and after eating) were used for analysis (Supplementary Table1). Subjects were returned to the colony room after the end of the recording session. The experiments were performed once a day for each subject.

### Marmoset marker-less 3D motion-tracking system

Our motion-tracking system software package for depth camera calibration, 3D data acquisition, and fundamental setup for physical simulation is available online (3DTracker-FAB, https://www.3dtrack.org). This motion-tracking system allowed us to robustly estimate the 3D trajectory of marmoset body parts as the positions of skeleton model parts (*Head, Neck, Trunk, Hip*) fitted by the physical simulation to the 3D point data of marmoset body shape (3D point cloud)(9)(34). The *Face* position was estimated by projecting the rectangle of the marmoset face area onto the 3D point cloud. This face area was detected frame-by-frame on 2D-RGB images by an object recognition algorithm YOLO3(35). To achieve sufficient accuracy, the YOLO3 detector was trained to detect marmoset face regions using 2,000 manually detected faces from two of the four marmosets (Marmo1 and Marmo2). 3D points inside the projected face rectangle were filtered to be less than 2.5 cm away from the center of the *Head*, and the average position of these 3D points was defined as the *Face* position (Fig.2c). Estimated error of *Face* position was reported in Supplementary Fig. 3.

### Behavioral data under chemogenetic neuronal manipulation

We used behavioral data obtained from an adult male marmoset that received a viral vector injection in the unilateral SN for expressing a Designer Receptor Exclusively Activated by a Designer Drug (DREADD) (AAV2.1-hSyn1-hM3Dq-IRES-AcGFP)(9). Using this DREADD setup, SN neurons are excited when the receptor is activated by administration of the agonist DCZ (HY-42110, MedChemExpress, NJ, USA; 3 μg/kg, orally). Two sets of free-moving behavior data (∼60 min) following either DCZ or vehicle alone (saline with 2.5% dimethyl sulfoxide FUJIFILM Wako Pure Chemical., Osaka, Japan) were used.

### Data preprocessing

All data preprocessing were performed using R version 4.0.3 (www.r-project.org) and its packages tidyverse version 1.3.0(45), data.table version 1.13.2, patchwork version 1.1.1, and magick version 2.7.0.

#### Marmosets

For free-feeding behavior analysis, the trajectory of body parts (*Face, Head, Trunk*, and *Hip*) was filtered with a locally estimated scatterplot smoothing filter using the stats::loess() function with span = 1/30 and downsampled to 10 Hz. Then, the spatial movement speed of each body part was calculated and the data coordinates were transformed frame-by-frame to make them posture-centered. Specifically, distance-from-center coordinates were transformed into posture coordinates with the *Trunk*’s center and the vector from *Trunk* to *Head* contained in the front-up quadrant. As a result, we used a set of 13 posture parameters (*Face* [x, y, z, v], *Head* [x ≡ 0, y, z, v], *Trunk* [x ≡ 0, y ≡ 0, z, v], and *Hip* [x, y, z, v], Fig. 2d). For SMP analysis, PC1 and PC2 were calculated using the stats::prcomp() function in R with scale = TRUE (Fig. 2e), and then scaled to a maximum absolute value of 1.

The same procedures were used for the data obtained after chemogenetic neuronal manipulation, except that instead of the *Face* position, we included the relative velocity of body parts into the SMP analysis, which was calculated from the postures at t+3 s (Fig. 5d) and normalized by the coordinates of the body at time t and the distances between these parts. Thus, the analysis of the neuronal manipulation data used a set of 26 parameters (*Head* [x, y, z ≡ 0], *Trunk* [x ≡0, y ≡ 0, z ≡0], *Hip* [x, y, z], their positions at t+3 s [x, y, z], their velocities [x, y, z, and absolute value]).

#### Macaques

For this analysis, we used published 3D macaque tracking data captured by OpenMonkeyStudio(14). From the published data, we extracted 3,534 s of data that was divided into 29 subsets with few missing frames (frame gap ≦ 20 frames) and sufficient duration (d ≧ 100 s). Then, the extracted data were transformed frame-by-frame from distance-from-center coordinates to posture coordinates with the *Hip*’s center and the vector from *Hip* to *Neck* contained in the front-up quadrant. These posture-coordinated parameters were interpolated in 2/3 Hz using the stats::loess() function with a span set to one-fifth of the data fragment length. As a result, we use 5,452 video frames with 36 posture parameters ([x, y, z] coordinates of 13 body keypoints with *Hip_x, Hip_z, Neck_z* scaled to zero, Fig. 4a). For SMP analysis, PC1 and PC2 extracted from these 36-parameter data were calculated using the stats::prcomp() function with scale = TRUE.

### Computational segmentation

The joint probability distribution of motion-motif length and class can be estimated by a blocked Gibbs sampler in which all motion motifs and their classes in the observed data (PC scores) are sampled. First, all data are randomly divided into motion motifs and classified. Next, motion motifs obtained by a part of the data are excluded from the dataset, and the model parameters are updated. By iterating this procedure, the parameters can be optimized. Parameter estimation in each iteration is described as forward filtering-backward sampling, which can be considered a maximum likelihood estimation process.

In the forward filtering step, the probability *α* that a data point given time step *t* is the ending point of a motion motif of length *k* classified motif class *c* was calculated as follows:

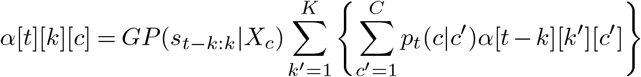

where *s*_*i*_ denotes time series PC score at a time step *i, C* denotes the maximum number of motion motif classes, *K* denotes the maximum length of motion motifs, and *p*_*t*_(*c*|*c*′) denotes posterior transition probability from motif *c*′ to *c*. The *GP*(*s*_*t*−*k*:*k*_|*X*_*c*_) is a predictive distribution of a Gaussian process of class c fitted to PC score *s* at time step from *t* − *k* to *k*, and is computed as follows:

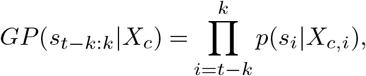

where *X*_*c*_ denotes a set of segments that are classified into class *c*. The variation within a segment is regressed on data fragments belonging to the same class *c* as follows:

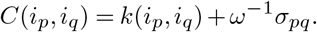

The class *c*_*i*_ of the *i*th motion motif is determined by the (*i* − 1)th segment and transition probability *π*_*c*_, which is generated from the *β*, generated Dirichlet process (DP) parameterized by *η* as follows:

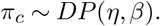

This *β* is an infinite-dimensional multinomial distribution, and its parameters were constructed by repeatedly breaking a stick, the length of which is one, with a ratio *v*_*k*_ sampled from a beta distribution, as follows:

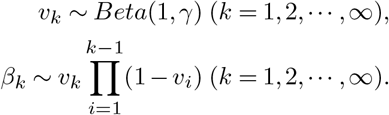

This stick-breaking process is also the DP, so the two-phase DP to generate *π*_*c*_ is called an HDP. In both the macaque and marmoset behavior analysis, the parameters were set *ω* = 10.0, *η* = 10.0, and *γ* = 1.0, and the length of the motion motifs was set around 30 frames (10 and 70, min and max, respectively). This model was reported as HDP-GP-HSMM for human motion analysis (29). For exploratory behavior analysis in NHP, we optimized the kernel function *k*(·, ·) with comparison as follows,

SMP and Model1, *k*(*i*_*p*_, *i*_*q*_) = exp(−||*i*_*p*_ − *i*_*q*_||^2^);

Model2, *k*(*i*_*p*_, *i*_*q*_) = 1 + *i*_*p*_ * *i*_*q*_; and Model3, *k*(*i*_*p*_, *i*_*q*_) = 1.

In addition, after learning the parameters for the data by iterating the model 100 times, the Viterbi algorithm estimated the most likely path of the latent state in the data. That is, all possible *α* in the data were obtained, and the most probable of all possible state sequences, combined with the learned transition probabilities, was adopted as the posterior state sequence.

### Posture model

For comparisons, the conventional posture model was estimated as described below. Standardized nine-dimensional marmoset motion information, the 3D coordinates of the four body parts (*Face, Head, Trunk*, and *Hip*), excluding the three scaled to zero, were mapped to two dimensions using the umap.UMAP() function in Python 3.741(46). Posture clusters were detected on the UMAP scores using the stats::kmeans() function in R. The representative posture for each cluster was the average of the data contained in each class.

### Synthesis of typical body motion sequences

The inverse operation of PC analysis was used to synthesize a typical posture series from the estimated PC coordinates. In each motif class, the typical PC waveforms (Fig. 2h, and Fig. 4c) were calculated within 10 Hz by moving the average for all PCs. These typical PCs were multiplied by the pseudo-inverse matrix of the eigenvector for the PC analysis of the test data that was calculated using MASS::ginv() function in R. Finally, the resulting values were inversely standardized using the mean and variance values of the data to synthesize the posture parameters. In the cases of marmoset free-feeding behavior and macaque ethogram detection, the postural information is displayed side-by-side along the time axis (Fig. 3c-e, Fig. 4d). The results of neural manipulation analysis visualized the motif’s internal variability by integrating the velocity relative to the initial position (Fig. 5g).

### Statistical analysis

Pearson’s Chi-square test, implemented with the stats::chisq.test() function in R, was used to compare intervals containing constant-specific timing among feeding subtypes and to test for differences in motion unit distributions among individuals. The Brunner-Munzel test, implemented with the lawstats::brunner.munzel.test() function in R, was used to compare the median value of segment length.

## ACKNOWLEDGEMENTS

We thank J. Kamei, R. Yamaguchi, Y. Matsuda, Y. Sugii, K. Yamashita, Y. Iwasawa, M. Nakano, and M. Fujiwara for their technical assistance. We also thank Drs. K. Shimatani (Institute of Statistical Mathematics), S. Nakamura (Tokyo University of Agriculture and Technology), and T. Taniguchi (Ritsumeikan University) for their comments on an earlier version of the manuscript.

This study was supported by MEXT/JSPS KAKENHI Grant Numbers JP17H06040, JP19H04996, 22K07338 (to KM), 22H05157 (to JM), JP17H02219 (to TH), and JP20H05955 (to TM), AMED under Grant Numbers JP20dm0107146 (to TM), JP20dm0307007 (to TH), JP20dm0207072 (to MH), and Takeda Science Foundation (to JM and HN).

## Notes

### Competing Interest Statement

The authors have declared no competing interest.

### Summary of Updates

update title

