## Supplemental Information for "Unsupervised decomposition of natural monkey behavior into a sequence of motion motifs"

### Supplementary Information

#### Supplementary Table 1. Summary of subject information and the feeding behavior test

| | Sex | Age (y) | Observed number | | | Test duration (s) |
|---|---|---|---|---|---|---|
| | | | floor-hand | floor-head | wall | |
| Marmo1 | F | 2.7 | 10 | 5 | 4 | 340 |
| Marmo2 | F | 1.4 | 5 | 2 | 3 | 200 |
| Marmo3 | M | 6.4 | 9 | 3 | 3 | 220 |
| Marmo4 | M | 5.1 | 7 | 5 | 3 | 260 |

**Supplementary Table 2: Distribution of observed motion motifs for each individual marmoset**

| | Motion unit | | | | | | | | | | | | | | | | | |
|---|---|---|---|---|---|---|---|---|---|---|---|---|---|---|---|---|---|---|
| | 1 | 2 | 3 | 4 | 5 | 6 | 7 | 8 | 9 | 10 | 11 | 12 | 13 | 14 | 15 | 16 | 17 | 18 |
| Marmo1 | 3 | 4 | 8 | 0 | 4 | 5 | 6 | 1 | 2 | 2 | 3 | 2 | 1 | 4 | 5 | 5 | 2 | 7 |
| Marmo2 | 10 | 5 | 6 | 3 | 7 | 8 | 10 | 6 | 3 | 0 | 7 | 5 | 6 | 4 | 2 | 1 | 3 | 1 |
| Marmo3 | 6 | 5 | 6 | 4 | 3 | 8 | 7 | 2 | 0 | 4 | 1 | 1 | 3 | 1 | 6 | 3 | 2 | 5 |
| Marmo4 | 1 | 8 | 7 | 1 | 9 | 6 | 12 | 3 | 3 | 2 | 5 | 6 | 1 | 7 | 8 | 8 | 0 | 7 |

**Supplementary Table 3. Distribution of posture clusters at feeding timing using the posture model (k = 18)**

| Feeding subtype | Posture cluster | | | | | | | | | | | | | | | | | |
|---|---|---|---|---|---|---|---|---|---|---|---|---|---|---|---|---|---|---|
| | 1 | 2 | 3 | 4 | 5 | 6 | 7 | 8 | 9 | 10 | 11 | 12 | 13 | 14 | 15 | 16 | 17 | 18 |
| wall | 8 | 4 | 1 | 0 | 0 | 0 | 0 | 0 | 0 | 0 | 0 | 0 | 0 | 0 | 0 | 0 | 0 | 0 |
| floor-head | 0 | 0 | 0 | 0 | 0 | 0 | 0 | 0 | 0 | 0 | 1 | 1 | 1 | 1 | 3 | 8 | 0 | 0 |
| floor-hand | 0 | 1 | 0 | 0 | 0 | 0 | 0 | 0 | 0 | 0 | 0 | 1 | 1 | 2 | 8 | 15 | 2 | 1 |

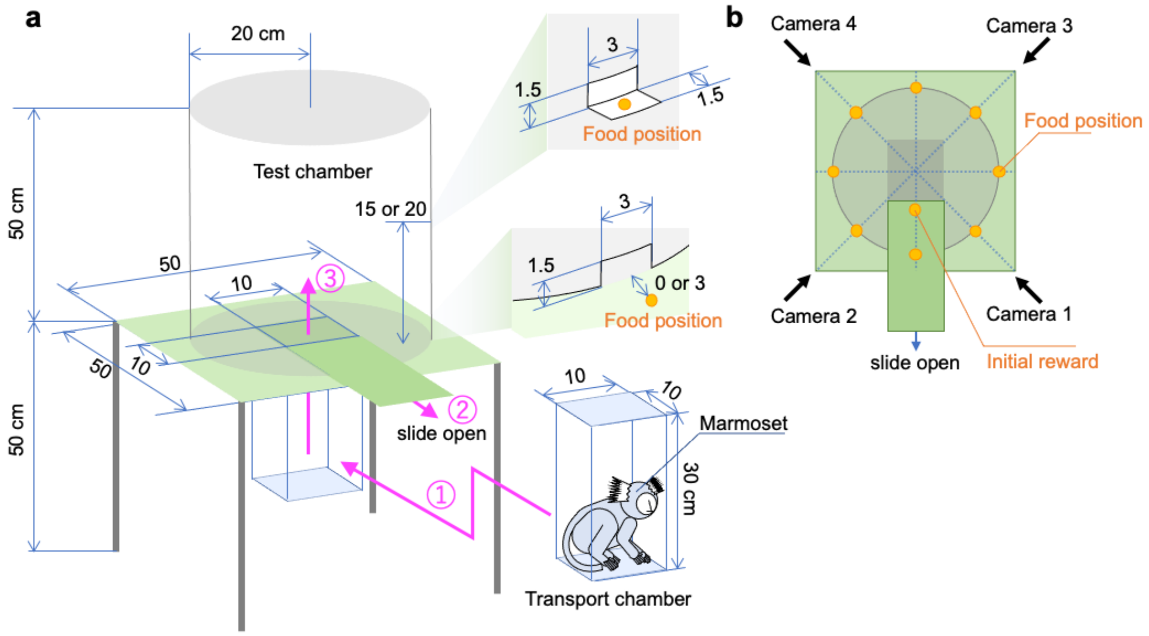


**Supplementary Figure 1 Experimental setup**

**(a)** Animals entered the behavior test chamber from the transport chamber by opening the door at the bottom of the test chamber floor. They then freely ate the food set in the small windows on the floor and the wall. A piece of food was placed at the entrance to guide the animals into the test chamber. **(b)** Four depth cameras were placed at 45-degree intervals to record three-dimensional behavior.

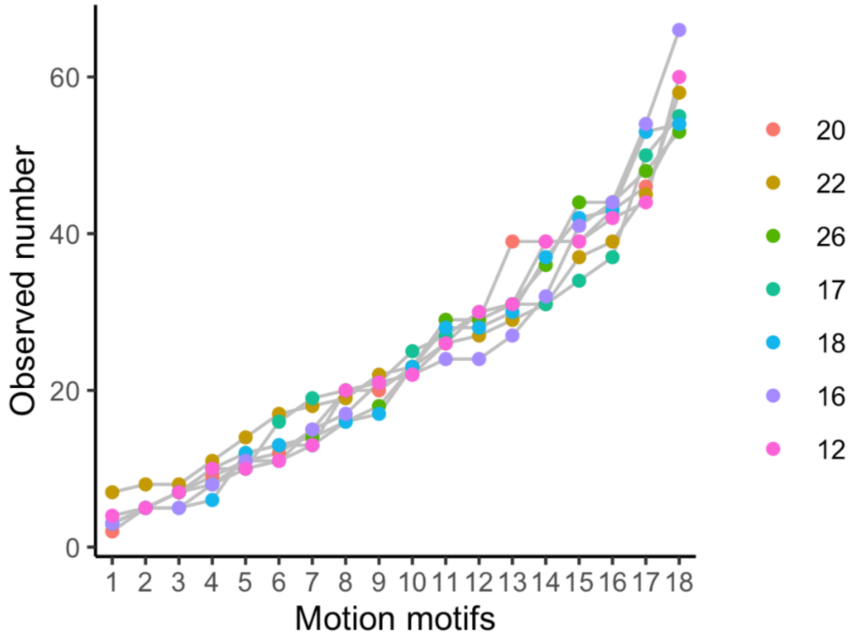


**Supplementary Figure 2 Distribution of observed segment number for each motion motif**

In the seven simulation results that yielded 18 class motion motifs, the frequency distribution of the motifs was similar regardless of the initial value of the class size (X-sq = 30.13, df = 102, p-value = 1.00, Chi-squared test).

a

b

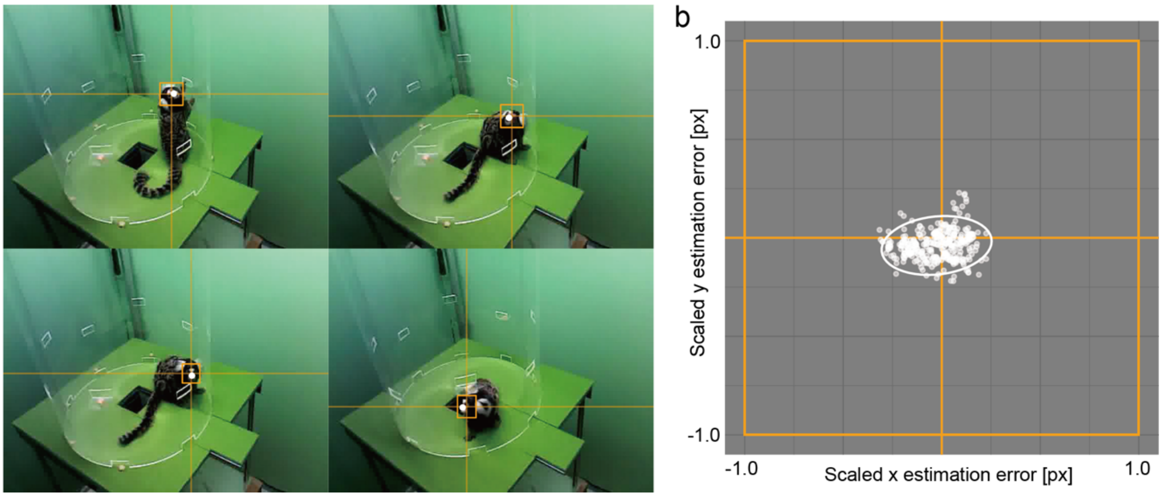


**Supplementary Figure 3 Estimation error of marmoset Face direction**

**(a)** Examples of 2D projection images of the *Face* position (orange rectangle with x and y axes) estimated by YOLO3 and those that were manually annotated (white point). **(b)** Scatter and density error plots between manual annotation and estimated results and 95% confidence ellipse on 2D images calibrated with the orange *Face* rectangle sizes in 350 randomly selected frames. Mean ± SEM errors were -0.026 ± 0.0066 4 and -0.037 ± 0.0039, on the X- and Y- axes, respectively.

**Descriptions of Additional Supplementary Files**

**Supplementary Movie 1**
**Description**: Free-feeding behavior of common marmoset were recorded by four depth cameras. Overlayed images with estimated body motion with four body parts, *Face*, *Head*, *Trunk*, and *Hip* and RGB camera image (left), and top and vertical projection of the trajectories (right).

**Supplementary Movie 2**
**Description**: Comparison of the marmoset free-feeding behavior and the synthesized behavior based on the estimated results. Body motion corresponding to the top left RGB video (bottom right), representative motion synthesized from the SMP motion motifs (middle bottom), and stop-motion-like animation synthesized by the conventional posture-clustering method with k = 6 (bottom right).

**Supplementary Movie 3**
**Description**: A set of 10 kinds of SMP motion motifs detected from freely moving macaque. Red dots representing the two body parts, *Nose* and *Head*. Horizontal position of *Hip* scaled to zero.

**Supplementary Movie 4**
**Description**: A set of 7 kinds of SMP motion motifs detected from freely moving marmosets with chemogenetic neural manipulation.
